# Curriculum-based outdoor learning for children aged 9-11: A qualitative analysis of pupils’ and teachers’ views

**DOI:** 10.1101/536441

**Authors:** Emily Marchant, Charlotte Todd, Roxanne Cooksey, Samuel Dredge, Hope Jones, David Reynolds, Gareth Stratton, Russell Dwyer, Ronan Lyons, Sinead Brophy

## Abstract

The relationship between child health, wellbeing and education demonstrates that healthier and happier children achieve higher educational attainment. An engaging curriculum that facilitates children in achieving their academic potential has strong implications for educational outcomes, future employment prospects and health and wellbeing during adulthood. Outdoor learning is a pedagogical approach used to enrich learning, enhance school engagement and improve pupil health and wellbeing. However, its non-traditional means of achieving curricular aims are not yet recognised beyond the early years by education inspectorates. This requires evidence into its acceptability from those at the forefront of delivery. This study aimed to explore headteachers’, teachers’ and pupils’ views and experiences of an outdoor learning programme within the key stage two curriculum (ages 9-11). We examine the process of implementation to offer case study evidence through 1:1 interviews with headteachers (n=3) and teachers (n=10) and focus groups with pupils aged 9-11 (n=10) from three primary schools. Interviews and focus groups were conducted at baseline and six months into implementation. Schools introduced regular outdoor learning within the curriculum. This study found a variety of perceived benefits for pupils and schools. Pupils and teachers noticed improvements in pupils’ engagement with learning, concentration and behaviour, as well as positive impacts on health and wellbeing and teachers’ job satisfaction. Curriculum demands including testing and evidencing work were barriers to implementation, in addition to safety concerns, resources and teacher confidence. Participants supported outdoor learning as a curriculum-based programme for older primary school pupils. However, embedding outdoor learning within the curriculum requires education inspectorates to place higher value on this approach in achieving curricular aims, alongside greater acknowledgment of the wider benefits to children which current measurements do not capture.

## Introduction

A mutual relationship between health, wellbeing and education exists. Evidence demonstrates that healthier children have higher educational attainment[1]. This association is mirrored, with research showing the social impact of education on health outcomes throughout the life course[1]. Thus, investing in a child’s learning has potential in maximising future achievement, employment prospects and health and wellbeing during adulthood. The school setting provides an opportunity to deliver a curriculum that engages children to reach their academic potential and define their future health outcomes and socio-economic pathway, reducing inequalities in health and education.

However, with schools currently facing a multitude of external, top-down pressures on educational attainment and health and wellbeing inequalities[2], there is a need for learning experiences that simultaneously improve health, wellbeing and school engagement whilst addressing curriculum needs. The opportunity to provide high-quality teaching experiences to engage children in learning is not solely restricted to the classroom setting. Taking learning outside the classroom and into the natural environment provides the opportunity for an integrated, cross-curricular approach to achieving education aims[3].

Outdoor learning encompasses a spectrum of curricular school activities that take place in the natural environment within school grounds or in the context of the local area. This ranges from broad nature-based learning such as Forest Schools, residential trips and outdoor adventure, to learning programmes tailored specifically to the core curriculum. This huge variation in the practice and understanding of outdoor learning means that the evidence base, whilst growing, shows huge variability in terms of the duration and type of outdoor learning offered, the target population involved and the outcome measures assessed[4].

In recent years, curriculum-based outdoor learning delivered by teachers in school grounds or the local area has gained momentum and is receiving attention from education experts and political figures alike[5]. This is particularly important as research suggests children’s wellbeing and mental health is declining and regular physical activity and engaging with the outdoors could potentially improve health, wellbeing and education outcomes[6–9]. It has been shown that delivering lessons in the outdoor environment can enrich learning and engagement, widen skill development and improve health, wellbeing and enjoyment in school[10]. However, despite its recognition at policy level, outdoor learning provision is still underutilised in primary schools, particularly beyond the early years[11]. Furthermore, research has demonstrated a marked decline in outdoor learning experiences between the early years and the later stages of primary education[12].

Efforts to integrate outdoor learning into the curriculum have been witnessed alongside curriculum reform across the United Kingdom[13]. In 2010, Wales introduced the Foundation Phase curriculum stage for ages 3-7, with a vision of encouraging ‘children to be creative and imaginative, and make learning more enjoyable and effective’[14]. This curriculum framework facilitates experiential learning through outdoor learning. However, despite government recognition of the benefits of outdoor learning in enhancing children’s social, physical, creative, cultural and personal development[15], there are no clear efforts to bring regular outdoor learning to older primary school age groups. In addition, conflict exists between the wider benefits to education attributed to outdoor learning, and the lack of measurement and the value placed upon these by education inspectorates.

As with many school interventions, the implementation of outdoor learning within the curriculum has not come without its challenges. Common barriers cited by teachers and headteachers include; existing curriculum pressure, the high demand on teachers’ time, teachers’ confidence and self-efficacy, safety, cost and access to resources and training[16–20]. Recommendations to overcome barriers and integrate outdoor learning within the school setting include providing schools with a clear evidence base[16].

Whilst research regarding the benefits of outdoor learning has examined cognitive, affective, interpersonal, social, physical health and behavioural impacts[21], there is a lack of research exploring the acceptability and mechanisms behind how outdoor learning can be effectively implemented on a regular basis by primary schools[22]. Furthermore, much of the literature aiming to gain the viewpoint of stakeholders has focussed solely on teachers and outdoor specialist staff [18, 19, 23, 24], highlighting the lack of experiences cited by pupils. If we are to create both meaningful education experiences in the outdoor environment, and ensure effective implementation of school-based programmes, it is essential to gain the viewpoint of not only those at the forefront of the delivery, but those who are recipients of such interventions, the pupils. Thus, the aim of this study was to examine the acceptability of an outdoor learning programme and to explore headteachers, teachers and pupils’ views and experiences of outdoor learning within the key stage two (KS2) curriculum (pupils aged 9-11). In addition, we examine the process of implementation to offer case study evidence to other schools who would like to offer outdoor learning to KS2 pupils.

## Methods

### Implementation

There was a general agreement among all schools that they intended to deliver at least one lesson outdoors a week. School A (the more urban of the three schools) chose to initially run outdoor learning in the school grounds but then became more involved with an outdoor activity provider utilising outdoor adventure as a key part of delivery as the project progressed. School B took a combined approach, initially delivered by a designated teacher trained in forest schools outside the school ground followed by teacher delivery. School C (the most rural of the three schools) took a teacher led approach utilising the local environment.

### Ethics

Ethical approval was granted by the College of Human and Health Sciences Research Ethics Committee (approval number 070117). All participants over the age of 18 (headteachers and teachers) provided informed written consent prior to participating. Pupils were required to provide informed written assent and parent consent in order to participate. All participants were reminded of their right to withdraw from the research at any point. All personal data such as names and school names was anonymised. Paper based data (consent) was stored securely in a locked cupboard and electronic data (interview transcripts) was stored in password protected documents on a secure University server.

### Approach

This study adopted a qualitative approach, viewed widely as the most suitable methodology in exploring barriers and facilitators of programme implementation[25]. Semi-structured interviews and focus groups were employed in order to gain an insight into the implementation of regular outdoor learning in the primary school setting. Interviews and focus groups are considered the most appropriate methods in examining the acceptability of interventions[26]. The process of thematic analysis generated themes and sub-themes from the data. The schools participating are members of the HAPPEN (Health & Attainment of Pupils in a Primary Education) Network), which aims to evaluate and share the evidence base for interventions currently delivered in primary schools, in order to improve children’s health, wellbeing and education outcomes[27]. The reporting of this study design is in accordance with the Consolidated Criteria for Reporting Qualitative Studies (COREQ)[28] (S1 Appendix).

### Participants

A convenience sample of three schools (School A, B and C) who expressed an interest in outdoor learning provision for their KS2 pupils were invited to take part in the research study. This sampling method was chosen to gather information-rich cases from schools committed to an outdoor learning programme[29]. Schools were contacted via a telephone conversation with the headteacher and were existing HAPPEN schools. The percentage of pupils eligible to receive free school meals at all three schools was below the national average (19%)[30]. Following headteacher consent, the lead researchers (EM and CT) presented about the research study at a school assembly to pupils aged 9 to 11 years (year 5 and 6 pupils) at each of the schools. Information sheets and consent/assent forms detailing the study aims were distributed to pupils, their parents and teachers within the school. Each assembly also provided the pupils and teachers an opportunity to ask questions related to the research project.

To participate in the research, children needed to provide written assent and parents needed to provide consent. Purposive sampling was used to recruit pupils for focus groups, ensuring an equal representation of age and gender. If any pupils were unavailable on the day, another person from this consented list was recruited. All three headteachers consented to take part in 1:1 interviews. Teachers from years five and six were invited to participate in a 1:1 interview. A purposive sample of consented teachers was selected to ensure an equal representation of gender.

### Data collection

This qualitative research study used focus groups with pupils at baseline (n=4) and follow up (n=6), 1:1 interviews with teachers (years 5 and 6) at baseline (n=4) and follow up (n=6), and 1:1 interviews with headteachers (n=3) at follow up. Interviews were conducted at two time points; baseline (beginning of intervention) (January 2017) and 6-month follow up (July 2017) in order to gather views at the start of the intervention and once outdoor learning was embedded within the curriculum. Interviews with headteachers and teachers were conducted according to individual preference, either by telephone or face to face on the school premises by one researcher (EM or CT). Pupil focus groups were completed within a private room at the school setting, with two researchers present (EM, CT, RC, SB, SD, HJ). Lead researchers were both female, trained to Masters level and had previous experience in conducting interviews and focus groups with both adults and children. Each focus group consisted of between six and eight pupils[31], aged 9-11 years of mixed genders. All interviews and focus groups followed a semi-structured topic guide (S2 Appendix), allowing deeper exploration of subjects including experience, views and opinions on outdoor learning, as well as suggestions for effective implementation in other schools. Applying open-ended questions to interviews allowed participants’ views to be explored further and topics to be discussed in a natural manner with the interviewer[32]. A lead researcher facilitated the interview process (CT or EM), whilst the other researcher (RC, SB, SD, HJ) provided technical support (digitally recording) and made field notes on key responses. These notes were verbally summarised to interviewees at the end of each interview in order to gain respondent validation[33]. In order to achieve neutrality, researchers reminded the participants at the start of interviews and focus groups that they remained impartial and of the study aims. Participants’ personal viewpoints were encouraged, and researchers emphasised that there were no right or wrong answers. Interviews lasted between 12 and 52 minutes overall (average length: pupil focus groups 30 minutes; teacher interviews 22 minutes; headteacher interviews 33 minutes).

### Data analysis

All interviews were digitally recorded and transcribed verbatim. Each transcript followed an open coding process by two researchers (EM, CT, SD, RC) independently and their responses were compared. Open coding allowed participants’ views to be summarized by assigning words or phrases to quotes. Codes were compared between researchers to ensure accuracy and consistency. If there was a discrepancy or disagreement in coding a third researcher adjudicated. All topics were compared with the written notes taken on the day of the focus group that had been agreed with the participants as an accurate account of their responses. Following this, two researchers worked together through an extensive process to discuss all codes and categorise them under theme and sub-theme headings (S3 Appendix).

## Results

Three key themes emerged from the transcripts; (1) Expectations and experience of outdoor learning, (2) Factors influencing outdoor learning and (3) Perceived impact on learning, health and development, all of which will be discussed in this section, alongside any suggestions in relation to each theme.

### Expectations and experience of outdoor learning

A prominent theme was the expectations and experiences of pupils and teachers regarding outdoor learning. This theme comprised of three sub-themes including feeling free, exposure to environment and safety and pupil engagement.

#### Feeling free

At baseline, pupils believed that outdoor learning would provide an escape from the uncomfortable and restricted conditions of the classroom. This escape from the classroom excited pupils, with discussions of freedom at both time points;

> *“So if you’re in a cramped classroom you don’t have that much room, if it’s wet play you don’t have that much room to do activities but if it’s outside you have loads of room”.* (Pupil, School B, Baseline)

Pupils also highlighted associations between fresh air, feeling more energised and an increased engagement with learning;

> *“And when we’re outside, like we get the fresh air, on a hot day if we’re in class we’re just boiling we won’t do as much work and we won’t do it as good, so when we’re outside we get fresh air”.* (Pupil, School B, Follow up)

Teachers believed the freedom allowed pupils to express themselves;

> *“I know lots of children that don’t cope very well with being in one classroom all day every day, they find it difficult to sit down but also for children who are more creative, they’ve got more opportunities to show that outdoors, I mean it’s the freedom and the movement and the expression and being able to use their bodies not just their voices and their hand, I mean it’s…Yeah, I think they all benefit”.* (Teacher, School B, Follow up)

In addition, outdoor learning offered pupils the ability to engage with play, an essential element of childhood;

> *“And it’s like really fun, because like the whole class goes out and if you’re like… most of the yard is by yourselves, because it’s kind of like playtime but you’re learning”.* (Pupil, School A, Baseline)

#### Exposure to environment and safety

Pupils suggested that the addition of outdoor learning to the curriculum would increase their exposure to the environment and their engagement with nature, expanding their learning;

> *“You learn about the outside world, you notice things about nature you never knew and you do different topics, but outside instead of inside”.* (Pupil, School A, Baseline)

This exposure to the natural environment was viewed as a positive aspect of outdoor learning during follow-up interviews, allowing pupils to learn about the outdoors. The opportunity to engage with nature at follow-up also encouraged an element of play;

> *“Because [being in the] woods like it’s more adventurous because you can just pick up sticks and start playing with them”.* (Pupil, School C, Follow up)

However, increased exposure to the environment was also felt to pose a risk to pupils and teachers regarding safety. At baseline, safety fears by pupils included physical injuries such as hurting themselves, or worries over getting lost from the rest of the class, something that the security of having physical boundaries in the classroom eliminated;

> *“You might hurt yourself on some bad things outside”.* (Pupil, School A, Baseline)

Despite perceived safety fears, pupils expressed frustration at the level of protection by teachers in the outdoor environment;

> *“That’s why a lot of people go off on that day because like the teachers are like really, they treat you like babies in the woods, they won’t even let you run”*. (Pupil, School B, Follow up)

Safety was initially a worry for teachers, however developing clear rules and boundaries and embedding outdoor learning into school life reduced the likelihood of any injuries;

> *“…initially there was things like trips and falls and head bumps and things like that and, touch wood, I’m not seeing so much of it so it’s embedded in the rules and things that we talk about. And when they climb the trees if it’s wet they’re only allowed up to an adult’s hip, if it’s dry they can go up to the shoulder and higher, they have to hold on, and there’s clear rules there and they really do stick to it, so”*. (Teacher, School C, Follow up)

#### Pupil engagement

Teachers discussed a range of ways outdoor learning engaged with all pupils;

> *“They love it, they absolutely love it, they just love it, yeah, there’s no problems. And they never, you know, have to walk with their heads down, you know, unhappy to go to the door, they’re always bouncing out to do something, so it’s just great, it’s really good”.* (Teacher, School A, Baseline)

In particular, outdoor learning engaged pupils of all abilities including those with behavioural difficulties and additional learning needs;

> *“They’ve [pupils] engaged in all activities that have been provided outdoors. So they definitely, it definitely engages all the children, whether they’ve got behavioural difficulties or not, I would say”.* (Teacher, School B, Baseline)

> *“So there are children who sit there very, very still and know how to, who know how they should behave socially or, you know, institutionalised, you know, they’re happy to do that, write neat, those kind of things that fit all those parameters, but for those children who don’t…I think that it’s more suited to them…It gives them, you know, an outlet and so yeah, I do think it’s for those children who learn perhaps in different ways, not just boys but yeah, but yeah probably boys”.* (Teacher, School A, Follow up)

The headteacher from this school also attributed the engagement by boys to the approach of outdoor learning;

> *“The teachers report as well how engaged they are, you know, with this style of learning and, you know, some of our perhaps more challenging boys particularly, you know, really enjoy the sort of the methodology, so yeah, I’ve heard nothing but positives”.* (Headteacher, School A, Follow up)

Engagement with learning was voiced by both teachers and headteachers, with a continuation of engagement during the follow up work in the classroom;

> *“I think it’s too much of a coincidence to say it’s not down to outdoor learning, because it’s an approach as well, you know, it’s not only the sessions outdoors, it’s what the sessions outdoors bring back into the classroom as well, isn’t it, and it’s the whole knock-on effect and it’s all about experience”.* (Headteacher, School C, Follow up)

In addition, how outdoor learning helped engage different styles of learners was discussed;

> *“You know, it’s not, you know without trying to be stereotypical, sometimes your very academic children they’re the ones that actually need it the most, because perhaps they’re quieter, they’re a little bit more book-based learners, the visual learners, so I think for those learners in particular, you know, so obviously you engage the learners who are kinaesthetic but also, you know, those other children, the ones that perhaps need it because potentially, you know, in the future they could be the ones who are, you know, in terms of looking after themselves and their wellbeing and so on, you’re perhaps hitting the mark with them and their sort of style of learning etc”*. (Headteacher, School A, Follow up)

Suggestions around increasing engagement and maintaining enthusiasm, related to ensuring the lessons conducted outdoors were fun and not more than once or twice a week, ensuring a novelty aspect.

> *“If we’re going to enjoy doing outdoor learning I think the lesson’s got to be fun, if it’s just like, I’m not sure, if it’s just like something boring and I’m not going to enjoy it as much and we’ll just start talking a bit”.* (Pupil, School A, Baseline)

> *“We’d get bored of it, I wouldn’t do every lesson, I think once or twice a week is enough, because otherwise, outdoor learning is supposed to be fun and if you keep doing it it can get a bit boring though”.* (Pupil, School A, Baseline)

### Factors influencing outdoor learning

Another theme to emerge from the transcripts encompassed the factors that influence the delivery of outdoor learning including motivations, curriculum pressure and accountability as well as natural and physical resources, support and teacher confidence.

#### Motivations

The implementation of outdoor learning was driven by headteachers and teachers’ motivations, including personal passion, passion of a colleague, pedagogical beliefs and a need to improve wellbeing outcomes. However, central to this subtheme was that of the rights of a child. Headteachers believed children had a right to be outdoors and that schools had a degree of responsibility in ensuring children were exposed to the outdoors in their learning;

> *“Every child is entitled, it’s their right to get outdoors and we have them all day, we have them for most of the daylight hours at certain times of year and so it’s our responsibility, I don’t think there’s a choice, I don’t think we can choose, shall we do it or shan’t we, we have to”. (Teacher, School B, Follow up).*

Other key motivations focused on improving pupil wellbeing and providing more opportunity to be outdoors;

> *“So, for me as well as their wellbeing, because I think that the children, there’s far too much time where children aren’t playing outside, they aren’t walking outside, they aren’t just outside, and I think a lot of that, with increasing volumes of children accessing counselling, spending a lot of time on social media, spending a lot of time on Xbox, a lot of time watching TV, they just don’t know the impact being outside has on their health and their wellbeing, and I’m really committed to developing pupils’ wellbeing”.* (Headteacher, School B, Follow up)

However, the high level of pressure placed upon schools by education inspectorates and the resulting resistance by the workforce was reinforced by one headteacher. This headteacher believed that in order to implement an initiative such as outdoor learning, an element of bravery was required by the school;

> *“And you know, you have pressures put on the school from Government, that goes down through the inspectorate, that passes onto the regional consortia, that’s passed onto schools, i.e. Headteachers, Governors, Senior Leaders, that’s passed onto the teachers, it’s passed onto the teaching assistants and it’s passed onto the pupils so it’s like a big pressure cooker and the whole system, you know, so until there’s that change in emphasis right at the top, you know, I think it will always be the brave schools that actually say ‘no, this is what I believe in and this is what we’ll do”’*. (Headteacher, School A, Follow up)

#### Curriculum pressure and accountability

The baseline interviews with teachers conveyed a feeling of overburden with some feeling that outdoor learning was an added pressure enforced by senior management at a time of high focus on academic literacy and numeracy targets;

> *“Until we’re up and running it seems like too much to do at the moment because all the emphasis is on literacy and numeracy all the time, that’s what the big push is at the moment and targets, so it just seems to be another new thing and another new pressure, despite the children enjoying going out”.* (Teacher, School A, Baseline)

Despite teachers generally feeling positive about outdoor learning, the academic pressures relating to evidencing work was at odds with the concept of teaching outdoors. This was particularly due to these teachers being responsible for a key stage that includes additional pressure and testing;

> *“Like the main concern for us, obviously, upper key stage 2 is obviously evidence of work, because there’s such a pressure now to have evidence, recorded evidence for every session or something in box, there’s a big pressure in that. …Again, lots of activities don’t provide evidence, so, it’s difficult then to gauge the amount of learning that they’ve done, apart from the bit of feedback basically”.* (Teacher, School B, Baseline)

Some teachers found it hard to design lessons with meaningful activities that could both encompass the concept of outdoor learning and meet the requirements of the curriculum.

> *“We’re at that early struggling stage looking for ideas of meaningful activities that we can do outdoors that do suit the outdoor environment and you’re not trying to directly lift a classroom activity into an outdoor activity, you’re trying to make it, you know, something that will work outdoors and there is a benefit, otherwise you might as well stay in the classroom and do it”*. (Teacher, School A, Baseline)

#### Natural resources

The schools included in this study had varied access to local natural environments, and this was acknowledged with reference to the types of lessons that were suited to this;

> *“Erm…. where our new school is and the surroundings we’ve got, obviously we’ve got access to the woodland area. We’re in a, you know, a really good spot that we can use, you know, we can use a lot more of it, it’s not just going outside, going into the yard, we can use the woodland which is great, you know, for Science, Geography-type lessons as wel”l.* (Teacher, School B, Baseline)

One headteacher highlighted that schools in a less fortunate position in terms of outdoor opportunities may struggle;

> *“In [city] lots of schools have aspirations to develop outdoor learning, but different schools have different challenges and different opportunities, isn’t it., other schools, perhaps who are in the middle of [city], number one, they don’t have woodland on their doorstep, so their opportunities to visit woodland would be limited”.* (Headteacher, School C, Follow up)

Indeed, utilising the immediate school grounds was raised as a challenge. One teacher at the more urban based school of the three felt that using the immediate school grounds was not enough for the older pupils, with the school later relying on external trips to provide pupils with an enhanced experience;

> *“Well, the stimulus is the trips, without the trips, as I, when I spoke to you the first time, you really struggle because you’re just using the school grounds, and lower down the school that’s not such a problem with building up their skills but by the time you get to the top end of the school, you need to branch out, you need to go further…But apart from identifying, we’ve got a little, we’ve got a small wooded area but apart from that it’s just grass really, so it was the trips that were the stimulus for, you know, all the extra ideas”.* (Teacher, School A, Follow up)

However, another school suggested relying on external trips would come at a cost, with parents having to fund the transport and staff needed to attend the trips. Teachers provided some suggestions to other schools;

> *“Prioritise anything that’s within walking distance of your school, so you know, if you have a river nearby or if you have a park nearby, that’s within walking distance, you know, utilise that as much as possible”* (Teacher, School A, Follow up)

#### Physical resources

In addition to the natural resources, the physical resources and time required to prepare new resources for outdoor learning were raised, with one teacher expressing their concerns over the transferability of traditional classroom lessons into the outdoor environment;

> *“Well because we don’t teach outdoors. We teach in the classroom, the things we do in the classroom, the resources we use are in the classroom and now we’ve got to, you either try and transfer those activities to an outdoor environment which is more challenging because of the resources, you know, the resources not being there, because everything is, we go more towards using computers now”.* (Teacher, School A, Baseline)

Another barrier highlighted by teachers was the clothing required for lessons, having to cancel if some children forgot coats etc. At follow up, one school had gained financial support, investing it in staffing and outdoor learning specific clothing;

> *“Supported financially, the school have bought waterproofs so that the weather’s not a barrier for the children and yes, they are funding me to continue in September for another year, so yes, very supported”.* (Teacher, School B, follow up)

Indeed, staff numbers were highlighted by schools as an obstacle to outdoor learning;

> *“Staff ratio, sometimes it, you know, when you want to do an activity you’d quite like it to be a group going out…we just haven’t got the staff sometimes to do these things or to go out, you know, it’s just it can be difficult to have the right staffing numbers out there”.* (Teacher, School A, follow up)

Funding was mentioned by all schools at follow-up. Improved access to funding resulted in resources moving from a barrier to outdoor learning to a facilitator;

> *“Like having ease of access to equipment has been another problem, so we’re trying to change that by we raised some money like I said doing this walk, trying to get equipment that can be accessed by the children and easily and not in a place where, you know, you need a member of staff to go with them”*. (Teacher, School A, Follow up)

The resourcing and funding was believed to be essential for ensuring a whole-school approach to outdoor learning;

> *“As I said, so I don’t want it to be a, sort of a flash in the pan so, you know, the resourcing element, the funding element, the fact it’s got to be a whole school sort of philosophy rather than turn to the whims of individuals”.* (Headteacher, School C, Follow up)

#### Support

The level of school, governor and parent support was highlighted by teachers and headteachers as an important factor. School B commented on the parental support throughout and how despite some initial concern and beliefs, general feedback and support from parents was positive;

> *“I did think we’d have a little bit of resistance at the beginning, because some parents believe children only learn by sitting at a desk, and indeed one grandparent did write on our Twitter account that, “A pity the children weren’t sitting at desks writing"…So, yes, the parents are very positive about the direction that we’re going, but it’s something that we will continue”*. (Headteacher, School B, Follow up)

Support from parents was also suggested by schools to overcome barriers associated with resources;

> *“..and when we were doing some outdoor activities we asked them to bring, you know, cardboard boxes and, you know, shelter making equipment and that kind of thing…So yeah, we did lean on parents somewhat”. (Teacher, School A, follow up)*

In addition to parents, support amongst the staff within school and utilising a whole-school approach was identified by one teacher as an essential element to effective implementation of outdoor learning;

> *“It’s obviously up to the school, you know, if they didn’t believe in it and they’re just going out for the sake of doing it, than I think it’s quite pointless then but if you are true believer in it and you can see value in it, I think you know you have to have your colleagues on board as well for it to work as a whole school initiative, if it’s something that you can do yourself, so long as you’ve got clarity as to why you’re doing it than I think it can work well really”.* (Teacher, School C, Follow up)

Teachers and headteachers commented on the support for outdoor learning by senior management and school governors, facilitated by communication between all levels of staff. Governor support was highlighted by all three headteachers as crucial, owing to the financial support, decision-making and strategic delivery that governors are responsible for.

> *“…So we wanted to involve all those partners within that model [of outdoor learning], and you know, that has come at a financial cost as well but the Governors were very committed and have released funds for that”.* (Headteacher, School A, Follow up)

Suggestions to overcome many of the resource and support difficulties experienced were often based upon shared practice both within and between schools;

> *“The Foundation [ages 4-7] first started it earlier than us originally so we, as a key stage 2, spoke to them, see what they did, went down to their classrooms and sort of spoke to them to see the kind of things they did. So I think it’s just communication as well isn’t it? Whether it, you know, whether it’s a meeting or a session or emails, just to see what everyone’s up to isn’t it, yeah”*. (Teacher, School B, Baseline)

With regards to between-school shared practice, School B also had a trained outdoor learning specialist and thus, were proactive in sharing their resources and providing training to other schools. The other two schools discussed a less formal approach, relying on sharing their experiences of outdoor learning with one another but these schools advocated for more shared practice and resources to aid implementation.

#### Teacher influence

Both pupils and teachers made links between the personality of a teacher and their enthusiasm with delivering outdoor learning. At baseline, teachers had mixed opinions of both their own and their colleagues’ confidence to deliver outdoor learning. Some felt a lack of knowledge left them in a position of low confidence, whilst others felt more confident in their ability to adapt lessons to the outdoor environment;

> *“As I say not at the moment, not personally…If I knew what I was doing yes but it’s coming up with the ideas in the first place, so I guess not”.* (Teacher, School A, Baseline)

> *“I think it’s brilliant, I feel confident that I can do it, I feel enthusiastic about it, I think it’s great for children to be given that freedom of being outside, and doing something which is going to help their learning, just because I think children find it difficult to be stuck behind a desk for so much of the time…Yeah, I feel quite confident, you know, obviously I’m looking around now, you know, what people have done, you know, in other schools that have worked…but yeah, I feel confident that you know, any lesson, if you said, can you put an outdoor slant on it”.* (Teacher, School A, Baseline)

Teacher confidence was also influenced by the expected workload and traditional learning approaches associated with this key stage;

> *“Right, initially, I thought, “Oh, no!”, because it’s upper school, you tend to focus a lot of written work in class, and obviously foundation phase are used to doing it, so it was a case of, “Oh, where do I start?”, initially. That was my first thought*… *No, I feel more confident now, now that it’s sort of implemented into my teaching. I do feel a bit more confident in preparing outdoor resources*…*I think teachers at foundation phase are a little bit more confident, and they tend to use, they’re doing it a lot more than we do”.* (Teacher, School B Baseline)

In addition, one teacher felt that for colleagues to buy into outdoor learning and feel confident to deliver the programme, it was important for learning objectives to be clear;

> *“…so long as they can see a point to outdoor learning, because there was a big myth when it started that we were just going to go out to the woods and play and it was going to be a free for all and I think that was the bit where they were saying “Oh what’s the point in this”, not just using it as a PE lesson, if they can see that there is a learning objective to it then I think it’s much more”.* (Teacher, School C, Follow up)

Those that had received prior training in formal outdoor programmes such as Forest Schools expressed higher confidence levels in delivering outdoor learning compared to those with less training;

> *“I’m quite confident myself, I’ve been forest school trained so it’s something that I’m more confident… I think we’ve had a lot of training now with it and the more we do it, obviously the more confident we get so”.* (Teacher, School C, Baseline)

The associations between consistent training, access to resources and teacher confidence was alluded to by a headteacher from another school, with this confidence impacting on how much outdoor learning was delivered at ground level. One headteacher also commented on the increase in confidence they had witnessed as the programme developed, indicating that increased experience in delivery resulted in higher levels of confidence;

> *“The other then is the confidence where, (name of teacher) has led, from just being apprehensive about taking children up to the woods, which is on our doorstep, as you know, all of a sudden he’s walking children on a five mile walk…you know, where that’s, in the past, a similar trip, we’d have had to pay for a guide to do that, he has the full confidence”.* (Headteacher, School C, Follow up)

### Perceived impact on learning & development

The perceived impact of outdoor learning on pupils’ learning and development emerged as a theme in relation to behaviour, concentration and memory, skill development and benefits to health and wellbeing.

#### Behaviour

There were mixed responses regarding the perceived impact on pupils’ behaviour. Those that believed it would have a negative effect at baseline made particular reference to the excitement of outdoor learning fuelling disruptive behaviour;

> *“If we were out, maybe like more start being hyper, because in class we probably have got discipline, once we start getting out and it’ll be exciting”*. (Pupil, School B, Baseline)

In comparison, other pupils felt that outdoor learning could improve behaviour through increased access to space;

> *“I think it’ll change our [behaviour], like if we do once or twice a week, then it would change our behaviour in a way, inside school like, so outside we’re not like fidgety, if we’re outside, it’s better”*. (Pupil, School B, Baseline)

At the start of the programme, teachers recognised an improvement in classroom behaviour and even an improvement in the quality of work produced by pupils;

> *“Yeah, because when you come indoors they’ve had their fresh air, they’re more likely to come in and you have that calm down time..and you get the better behaviour because they’ve had that chance to go …when they’ve been out, it’s just so much more, there’s just a better working atmosphere, so when you… the more opportunities to get out and about, up and down, not just doing something at a desk, the more quality work you’re going to get from them when you do ask them to sit at their desk I think”. (Teacher, School A, Baseline)*

This was also discussed in follow-up focus groups, with pupils making references to the effects of outdoor learning on subsequent behaviour in the school day;

> *“I kind of think it’s better with outside, but then when you go inside for class, or everyone’s a bit more tired so the, say not like running around as much in the class”.* (Pupil, School C, Follow up)

From the teachers’ perspective, improved behaviour and engagement with learning was displayed by children with additional learning needs and behavioural difficulties;

> *“We do have children that have challenging behaviour, but we find they are far more engaged outdoors than indoors”.* (Headteacher, School B, Follow up)

Indeed, while improved behaviour was voiced by all schools, particularly with regards to follow up work, others also believed behaviour was better during outdoor learning than in classroom-based lessons;

> *“No, we’ve got quite clear boundaries for them as well so it’s not sort of a case of we go down the woods and it’s a free for all, there’s very strict rules as to behaviour in the woods, they stick to the… in fact, I’d say they stick to rules better when we’re outdoors than they do when we’re inside but I think it does, you know, you can see the impact back in the classroom then after we’ve been, definitely”.* (Teacher, School C, Follow up)

The headteacher of this school recommended less affluent schools utilise pupil deprivation grants for outdoor learning as a suggestion to improve pupil behaviour;

> *“So, you know, I would urge, if I was a headteacher in one of those schools, I would…and the big deprivation grant, and if I was headteacher there now I would definitely look to utilise some of that deprivation grant to encourage outdoor learning, and I’m sure it would have a positive impact on pupil behaviour. And the thing is, it’s a vicious circle, isn’t it, if children aren’t behaving, they’re not learning”*. (Headteacher, School C, Follow up)

#### Concentration and memory

Pupils suggested at baseline that the introduction of outdoor learning within their school day would have an impact on their concentration and memory. From a positive perspective, this was discussed in relation to the feelings of comfort experienced by pupils;

> *“Yes, because when you’re outside you’re not all sweaty and you like can’t really concentrate that much when you’re like really sweaty but if you’re like outside you’re like nice and cool so it’ll help you listen better and concentrate better”.* (Pupil, School B, Baseline)

However, the impact of distractions on concentration was also brought up during follow up focus groups, whereby pupils commented on greater distractions outside. Indeed, whether being outdoors had a positive or negative effect on their concentration was debated among pupils and even internally by one pupil who demonstrated both sides of the argument;

> *“I think it does [improve concentration] but then it doesn’t because it like helps people get more excited and it makes you listen more but then also it doesn’t because we’re all talking all the time outside and it’s a lot louder so a lot of the time we don’t listen to what the teacher says”*. (Pupil, School A, Follow up)

> *“You could get distracted by cute dogs walking past, you could get distracted by trees blowing, you could get distracted by say if another child or pupil or class has been let out to play early, get distracted by them, a netball match or anything like that, you can just easily get distracted outside rather than in the classroom”*. (Pupil, School B, Baseline)

Pupils referred to the perceived effects on concentration in terms of the novelty of outdoor learning and the excitement and enthusiasm it would bring to pupils;

> *“It’s made us a lot more concentrated, like before in the old school we’d just be sitting there like, but now we’re like, concentrated, before we’d be like, oh, and it was really boring”.* (Pupil, School B, Follow up)

The increased space offered by learning outdoors was discussed by teachers who believed that this made pupils more focused on their learning;

> *“The only difficulty is I suppose is that sort of making your voice travel, and keeping them focused, but then you know, in class that there’s as much trouble there keeping them focused, because they’re sat close to each other on the carpet, you know, poking each other and stuff like that…I think if it’s clear, they go out, they’re focused on the task, they’ve got their own space to do it in, they’re not looking around, they’re not looking for distractions, they’re quite focused on what they’re doing, just because they’re outside doing”.* (Teacher, School A, Baseline)

#### Key skills development

Pupils and teachers discussed the range of skills that they could develop through engagement with outdoor learning, including communication and teamwork;

> *“I think that like it makes us like learn how to work as a team”.* (Pupil, School C, Follow up)

> *“They were much more able to collaborate outside as it’s kind of freedom of the class, they might work in different groups and, you know, you’re not expecting them, they share more easily”.* (Teacher, School A, Follow up)

A range of other skills were discussed by teachers, including problem-solving, discussion skills and independence skills.

> *“The opportunity to work as a group, you know, they love the activities, and they get challenge activities, so they’ve got to use their problem solving skills, they’ve got discussion skills”.* (Teacher, School B, Baseline)

Aside from learning specific skills, one headteacher believed that outdoor learning ensured children developed in a holistic way;

> *“Because it develops the whole child and it enables all of the children to develop those skills that children just don’t seem to have. For us, we see children that haven’t got the resilience, especially Year 6 children, they don’t have the resilience to deal with such normal childhood situations and matters because they haven’t interacted enough, they haven’t risk-taked…so we want the children to develop, the whole child, the ability to be good citizens, but if they’ve never worked in teams, if they’ve never lost, if they’ve never failed, they haven’t got those resilience, and then they haven’t got the perseverance then to keep on trying, keep on trying, keep on trying”*. (Headteacher, School B, Follow up)

#### Health and wellbeing

During the interviews, there was a feeling among pupils that an increase in utilisation of the outdoors would help to increase physical activity and fitness. Outdoor learning was seen as a means of providing an opportunity to reduce sedentary time associated with traditional classroom based lessons:

> *“Without going outside you can’t really keep fit and like indoors we’re pretty much just sitting down at a desk writing”.* (Pupil, School B, Follow up)

Indeed, many pupils advocated for increased opportunities to be active during outdoor learning lessons;

> *“More exercise and like maybe more games because what we did was looking around and just marking things off”.* (Pupil, School A, Follow up)

This included opportunities for increased physical activity as well as the addition of structured sports;

> *“If we were doing sports with it, not so much learning, but like sports as well learning”*. (Pupil, School B, Follow up)

Alongside the physical health benefits, pupils remarked upon the emotional health benefits in terms of a feeling of happiness and how this had a knock on effect with willingness to attend school;

> *“Yes, less bored and I think it’s much happier to go to school”*. (Pupil, School B, Follow up)

The discussion around wellbeing centred on the reduced sense of stress resulting from pupils learning outdoors;

> *“I think that they like us being outdoors because maybe they don’t like us feeling stressed because we could be stressed in the classroom and instead of being stressed we’re outdoors and we’re happy”. (Pupil, School B, follow up)*

This stress reduction was not limited to children, with one teacher also commenting to feel less stressed as a result of the outdoor learning programme;

> *“And I just think it’s, yeah, I think it’s stress relieving for teachers as well as children*”. (Teacher, School A, follow up)

Indeed, for a few teachers, the introduction and responsibility of delivering outdoor learning provided them with a sense of increased personal wellbeing and in particular, job satisfaction at a time of extreme pressure;

> *“Just that happy that it’s happening really… felt like a breath of fresh air and there …, being told by management and the head, let’s get outdoors, it’s like feeling like someone’s taken the shackles off us and oppressive feeling, so it have felt like a bit of fresh air around the school and there’s a new buzz…my feeling is just like, wow, this is just what I came into teaching for, this feels like teaching, whereas before it didn’t feel like teaching to me, it felt like Orwellian nightmare [laughs]”.* (Teacher, School A, Baseline)

## Discussion

The overall positive and enjoyable experience of outdoor learning reported by children in this study is echoed by a high number of studies reporting children’s experience of the outdoors[24,34,35]. Pupils described how outdoor learning provided them with feelings of freedom and fun and discussed this in relation to an escape from the restricted, physical environment of the classroom. This also provided the opportunity for pupils to engage in and learn through play. This sense of freedom is reinforced in some of the earlier literature on outdoor learning, in which one of the main advantages of using the outdoor environment was the ability for children to learn through moving freely and play[36]. This freedom of the outdoors also provides children with important multisensory experiences that contributes towards improvements in motor development[37] and motor and sensory stimulation[38].

Pupils and teachers in this study commented on increased engagement with learning in the outdoors and overall school engagement. Research has demonstrated the ability of the natural environment to promote a desire to learn[35] and a positive relationship between learning and school motivation[39]. Teachers in our study suggested pupils’ learning was facilitated through the experiential pedagogy of outdoor learning. Greater pupil engagement is reinforced in the literature in relation to experiential learning and the different pedagogy of outdoor learning, such as less confined outdoor spaces and outdoor resources[18].

Whilst the positives of the environment and exposure to nature were discussed, safety was initially a concern by both pupils and teachers and has been mirrored in other outdoor learning studies[40,41]. However, whilst some pupils were concerned over safety in our study, feeling restricted by teachers was a negative by others. Research into teachers’ pedagogical practice outside the classroom found that teachers’ fears of class control outdoors triggers increased authoritative teaching practices[42]. Indeed, many pupils may thrive over the physical and risk taking challenges the outdoor environment offers[43] and removing all elements of risk may remove the fun aspect reported by pupils. Once outdoor learning was embedded in this study, teachers did not report any incidents and felt safety was less of a concern as children were more aware of boundaries. The need for an initial adjustment period has been raised in the literature, whereby once outdoor learning became embedded and students adjusted to the different learning environment, discipline became less of an issue and the rewards more apparent[19]. For the effective implementation of outdoor learning, it is essential for schools to consider the balance of risk and benefit in relation to perceived safety fears and opportunities for outdoor play.

The notion felt by headteachers in this study that children have the right to be outdoors is supported by others[44]. The United Nations Convention on the Rights of the Child (UNCRC) movement[45] within schools has improved the understanding and application of children’s rights in recent years[46]. Teachers also felt that children have lost access to outdoor play environments. Indeed, the number of children participating in unstructured, outdoor play is decreasing[47]. With this in mind, outdoor play through outdoor learning may be one of the only opportunities children have to experience the natural environment[21,48]. This engagement with and exposure to nature was cited as a benefit by pupils and teachers. At a time when environmental issues and sustainability are high on both the education and political agenda, outdoor learning provides the opportunity to encourage children to become environmentally aware.

A key point of discussion by teachers was curriculum factors and accountability. This is unsurprising given the large amount of research showing curriculum pressure as a barrier to delivery of interventions in the school setting[2,49]. In relation to outdoor learning, research suggests that teachers’ values may be influenced by top-down, external curricular pressure, suggesting incongruity exists between the narrow measurements children are judged on and the wider aims of education[18]. In this study, teachers discussed feeling overburdened and initially viewing outdoor learning as an additional pressure. For outdoor learning to be successful, schools need to value it as a means of achieving curricular goals, not merely an add-on initiative or an activity in isolation to their teaching[22]. Indeed, research with teachers has suggested a clear focus on curriculum related benefits would encourage a higher uptake of outdoor learning[50]. Conversely, it is essential for education inspectorates to view and support outdoor learning as a method in achieving curricular aims and this should be mirrored in testing requirements in which schools are judged.

In this study, teachers highlighted the barrier of evidencing work in the outdoors. Possible methods to overcome this have been suggested including taking pictures of work conducted outdoors and asking children to annotate this, advocating for more shared practice with regards to methods of evidencing work done outdoors[34]. A report by the Welsh Education Inspectorate (ESTYN)[5] evaluating outdoor learning in Foundation Phase concluded that teachers assessed children’s learning ‘less often’ and ‘less well’ outdoors than in the classroom, allowing for important developmental milestones to be missed. With the current focus by education inspectorates on academic targets, particularly in the higher key stages, it is essential that educators develop appropriate methods and tools to assess these skill developments in line with curriculum testing requirements in order to find value in the outdoors as a setting for learning.

All schools in this study referred to the need for financial support. However, a report by Natural England stated that simply providing funding for outdoor learning activities was not the answer to increasing education outside the classroom, with many schools on low budgets demonstrating excellent practice in outdoor learning[50]. In addition to financial support, teachers in this study highlighted the importance of senior leadership and governor support and advocated for a whole-school approach through all levels of school staff. Research has demonstrated that senior staff support was a strong enabler for the uptake of outdoor learning, in addition to passionate, committed and enthusiastic teaching staff[50].

Teacher confidence as a barrier to outdoor learning was identified by teachers in this study and has been cited in previous research[21]. Teachers are considered agents of change in delivering school-based programmes[51] and factors such as teacher confidence and level of training can influence the delivery of these programmes[52]. Developing teacher confidence requires school-based, outdoor learning specific continuing professional development (CPD) training[16]. However, research into teacher CPD demonstrates that it takes about 30 hours of training to make a significant change in pedagogy[53]. This level of contact required with teachers is unlikely to be feasible within the scope of inset and training days, and given the current high demand on teacher workload. A longer term solution would be to provide more focus on outdoor learning specific training for older aged children in teacher training, though this would not support current teaching staff in need of development and training.

Perceived improvements in concentration highlighted by both pupils and teachers in this study is supported by research on the role of the natural environment and concentration using ‘attention restoration theory’[54,55]. This theory suggests mental fatigue and concentration can be improved through the effective restorative environment of the outdoors. Improvements in behaviour were also cited by teachers, particularly the ability of outdoor learning to engage pupils with behavioural difficulties or additional learning needs. In addition, pupils and teachers commented on the positive impact on key skill development such as interpersonal and social skills and the enhancement of relationships through teamwork, all of which are recognised in the literature[21]. However, pupils discussed the potential for distractions when working outdoors. Indeed, the outdoor environment transfers learning to a different learning space, one that requires children to balance their learning with background noise and distractions caused by the natural environment.

Pupils and teachers also discussed the health and wellbeing benefits of outdoor learning. Becker (2017)[6] highlighted physical activity (PA) and mental health as understudied outcomes. A benefit voiced by pupils in this study was the opportunity to be physically active and the reduction in time spent being sedentary. Pupils advocated for more opportunities to be active in outdoor lessons. With research demonstrating higher levels of PA being exhibited on outdoor learning days[56] and current upwards trends in sedentary behaviour, providing opportunities to be physically active during outdoor learning sessions could contribute to children’s overall physical activity.

Improvements in both pupil and teacher wellbeing were also highlighted in this study. Teachers reported feelings of increased job satisfaction and wellbeing, a finding that is mirrored in the literature[57]. Teacher wellbeing is considered a critical factor in creating a stable environment for pupils to learn[58] and has been associated with academic achievement[59]. However, much of the discourse around teacher wellbeing has focussed on the reported stress, burnout, workload and decline in teacher retention in recent years[60,61]. With this in mind, the benefits to teacher wellbeing and increased job satisfaction cited in this study suggest that outdoor learning may provide an avenue in fostering teacher wellbeing and creating learning contexts for pupils to succeed. Research has demonstrated that exposure to the natural environment in primary school plays a significant role in improving positive mental health and wellbeing for pupils[62,63]. Results from a recent systematic review also demonstrated the importance of access to green space on child mental wellbeing, overall health and cognitive development[64]. With research highlighting the relationship between health, wellbeing and education outcomes[1], results from this study highlight the potential for outdoor learning as a means of improving the health, wellbeing and education outcomes for children.

### Strengths and limitations

Findings from this study explore detailed experiences of outdoor learning from those at the forefront of delivery and implementation, headteachers and teachers. In particular, this paper contributes to the gap in experiences reported directly by pupils. The knowledge gained through interviews and focus groups from a whole-school perspective provides an opportunity for schools to reflect on the facilitators and potential challenges of implementing outdoor learning. This understanding of the barriers that schools have experienced encourages prospective schools to design and deliver tailored outdoor learning programmes.

There are a number of limitations to review when considering the findings from this study. The schools participating in this study all had a percentage of pupils eligible to receive free school meals below the national average (19%) and thus would be considered less deprived. Another limitation is the small sample included in this study, in particular the homogeneity of the schools and participants in relation to ethnicity. This may limit the transferability of the findings and requires future research to include larger sample sizes of socio-economic, ethnically, culturally and geographically broader populations. The schools included in this study all had access to green space or the natural environment within close proximity to the school setting. However, access to and availability of the natural environment was not recorded in this study. It is important for future research to explore the experiences and implementation processes of schools with limited access to the natural outdoor settings. In addition, research into the investment of school grounds to increase green space would be welcomed, thus bringing nature to schools. Despite these limitations, this study contributes towards the understanding of barriers and facilitators of an outdoor learning programme within the primary school curriculum. These findings provide schools committed to implementing outdoor learning with case study examples to ensure effective implementation to improve the health, wellbeing and education outcomes of pupils. Further research involving quantitative assessments of health, wellbeing and education outcomes would strengthen the knowledge base for schools and education inspectorates.

## Conclusions

Participants in this study supported the case for outdoor learning in the KS2 curriculum, identifying benefits ranging across the personal, social, physical and curricular domains. The schools in this study reported a variety of benefits of outdoor learning for both the child and the teacher and for improving health, wellbeing, education and engagement in school. Findings highlight that outdoor learning has the ability to enthuse, engage and support children of all learning abilities in reaching curricular aims alongside positive improvements to health and wellbeing. With the relationship between education and health well documented throughout the life course, this study supports outdoor learning as a method of facilitating pupils in achieving their academic potential, improving educational experiences and attainment and ultimately improving future health outcomes and employment pathways.

Importantly, this study contributes to the gap in experiences reported by both pupils and teachers of outdoor learning programmes in the older ages of primary schools. Findings from this study offer schools important insights into the barriers and facilitators of implementing a regular outdoor learning programme within the KS2 curriculum. However, these findings highlight the gap that exists between the health, wellbeing and wider educational benefits achieved through outdoor learning, the lack of tools in evidencing these and the narrow measurements in which schools are judged on by education inspectorates. Results from this study advocate for additional help and support from education inspectorates to enable schools to feel that ‘non-traditional’ learning methods are valued and can address the curriculum pressures in which schools are measured on. More support, training and engagement for schools as well as direction from inspectorates is required if outdoor learning is to become a more mainstream method in addressing curriculum aims.

## Author Contributions

Conceived and designed the study: EM, CT, SB. Conducted data collection: EM, CT, SB, RC, SD, HJ. Conducted data analysis: EM, CT, RC, SD. Wrote the paper: EM, CT. Provided critical input: DR, GS, RD. Provided supervision: SB, RL. RD acts as a school advisor to the HAPPEN project.

## Acknowledgments

The authors would like to thank all participating schools, headteachers, teachers and pupils that took part in this study. This work was supported by the National Centre for Population Health and Wellbeing Research.

